# A Highly Conserved 3_10_-Helix Within the Kinesin Motor Domain is Critical for Kinesin Function and Human Health

**DOI:** 10.1101/2020.09.22.308320

**Authors:** Aileen J. Lam, Lu Rao, Yuzu Anazawa, Kyoko Okada, Kyoko Chiba, Shinsuke Niwa, Arne Gennerich, Dan W. Nowakowski, Richard J. McKenney

## Abstract

KIF1A, a kinesin-3 family member, plays critical roles as a long-distance cargo-transporter within neurons. Over 100 known KIF1A mutations in humans result in *KIF1A* Associated Neurological Disease (KAND), developmental and degenerative neurological conditions for which there is no cure. A de novo missense mutation, P305L, was recently identified in several children diagnosed with KAND, but the underlying molecular basis for the disease phenotype is unknown. Interestingly, this residue is highly conserved in kinesin-family proteins, and together with adjacent conserved residues also implicated in KAND, forms an unusual 3_10_-helical element immediately C-terminal to loop-12 (L12, also known as the K-loop in KIF1A) in the conserved kinesin motor core. In KIF1A, the disordered K-loop contains a highly charged insertion of lysines that is thought to endow the motor with a high microtubule-association rate. Here, we characterize the molecular defects of the P305L mutation in KIF1A using genetic, biochemical, and single-molecule approaches. We find the mutation negatively impacts the velocity, run-length, and force generation of the motor. However, a much more dramatic effect is observed on the microtubule-association rate of the motor, revealing that the presence of an intact K-loop is not sufficient for its function. We hypothesize that an elusive K-loop conformation, mediated by formation of a highly conserved adjacent 3_10_-helix that is modulated via P305, is critically important for the kinesin-microtubule interaction. Importantly, we find that the function of this proline is conserved in the canonical kinesin, KIF5, revealing a fundamental principle of the kinesin motor mechanism.

## Introduction

Kinesins are molecular motor proteins that hydrolyze ATP to move processively along, or actively remodel, microtubules (MTs) within cells. The kinesin superfamily (KIF, (1)) in humans is large, containing ∼45 genes with diverse functions. KIF1A is a member of the large kinesin-3 (KIF1) sub-family known to participate in long-distance transport of various cellular cargos, primarily within neurons (2). KIF1A has unique biophysical properties, being among the fastest and most processive kinesins reported to date (2-5). The kinesin motor core is highly conserved among different families, with distinct insertions in loop regions often accounting for specific biophysical adaptations. KIF1A contains an elongated loop 12 (L12), within its motor core, often referred to as the ‘K-loop’ due to the presence of multiple tandem lysine residues, endowing it with a highly positive charge (6). The K-loop bestows KIF1A with an extremely high MT on-rate due to its presumed interaction with the highly negatively charged tubulin C-terminal tail domains (3, 7). The K-loop has never been visualized in any reported KIF1A structure, implying it is disordered in all states of the motor’s mechanochemical cycle observed thus far. Therefore, it is currently unclear if the K-loop must adopt a specific conformation or orientation, with respect to the MT surface, in order to effectively enhance the kinesin-MT association.

KIF1A has garnered much recent attention due to the fact that over 100 human mutations have been identified within the motor that lead to a group of developmental and degenerative neurological diseases now referred to as KIF1A Associated Neurological Disorder (KAND) (8). Within the spectrum of KAND phenotypes, many mutations were previously characterized to lead to phenotypes resembling Hereditary Spastic Paraplegia (HSP), a degenerative nervous system disorder resulting in progressive lower limb spasticity (9, 10). Interestingly, mutations in another transport kinesin, KIF5A, also result in HSP in humans (11). Given the large number of mutations mapped within KIF1A, it has become imperative to understand the molecular consequences of these alterations to the motor in efforts towards advancing a therapeutic strategy. The majority of KIF1A missense mutations map to the conserved motor domain (8), and several previous studies have characterized the molecular defects that result from some of these alterations (12-14). While the negative consequences of several mutations in highly conserved residues known to be important for kinesin’s enzymatic activity or MT-binding properties could be reasonably predicted, many mutations remain uncharacterized.

We recently found that several KIF1A mutations in the motor domain do not result in impaired motor motility, but rather result in overactive motors, presumably due to a disruption of KIF1A’s ill-defined autoinhibition mechanism (14). Given this finding, it is imperative to understand the molecular phenotypes of individual KIF1A mutations prior to any therapeutic attempt. Working with the grassroots patient advocacy group KIF1A.org, we recently became aware of a novel KIF1A missense mutation, P305L, which lies very close to the conserved K-loop in the motor domain. Patients with P305L mutation have developmental delay, cerebellar atrophy, ataxia and eye movement abnormalities, hallmarks of KAND (15). At the time, there was no molecular information about the role of this highly conserved proline residue in the kinesin motor mechanism, and its proximity to the K-loop implied that it could play a role in modulating the activity of this critical structural element. A recently published report (15) found that the mutation impaired KIF1A motor activity in cells and in multi-motor gliding assays, but a mechanistic understanding of the effects of this mutation on KIF1A motor activity is still lacking. There are now at least fourteen children diagnosed with this specific P305L mutation with substitutions by other amino acids not noted at this position (KIF1A.org, personal communication), indicating it is a common variant within the growing KAND population. Intriguingly, a mimetic mutation of the analogous residue in KIF5A (P278L) results in HSP in humans (16), suggesting this proline is part of a critical element for the conserved kinesin mechanochemical cycle.

The P305 residue within KIF1A is part of a highly conserved motif (PYRD/E), which we find to adopt a 3_10_-helical conformation in many families of kinesin motors from animals to fungi (see Fig. 2). 3_10_-helices differ from α-helices in the arrangement of their backbone hydrogen-bonding (with 10 atoms in the ring formed by making the H-bond), giving 3_10_-helices three, instead of 3.6, residues per turn. This arrangement results in tighter winding of the helix and repositioning of side-chains with respect to the more common α-helix. 3_10_-helices, which are more rare and are thought to be inherently more labile than α-helices (17-19), are typically short and have been proposed to act as intermediates in the folding/unfolding of alpha-helices (20). In addition, proline itself is an anomalous amino acid owing to its 5-membered ring within the polypeptide backbone, with ring-locked nitrogen unable to act as a hydrogen bond donor. Functionally, conserved prolines have been described to act as dynamic, flexible hinges that mediate propagation of conformational changes from one domain in a protein to another, such as in the activation mechanism of G-protein coupled receptors (21). And helix-distortion by prolines has been shown to mediate key mechanistic conformational changes, for example in the voltage gating mechanism of connexin32 (22). Thus, P305’s position at the N-terminal cap of the PYRD/E 3 10-helix may bestow this unique element with additional functional properties (see below).

**Figure 1.**
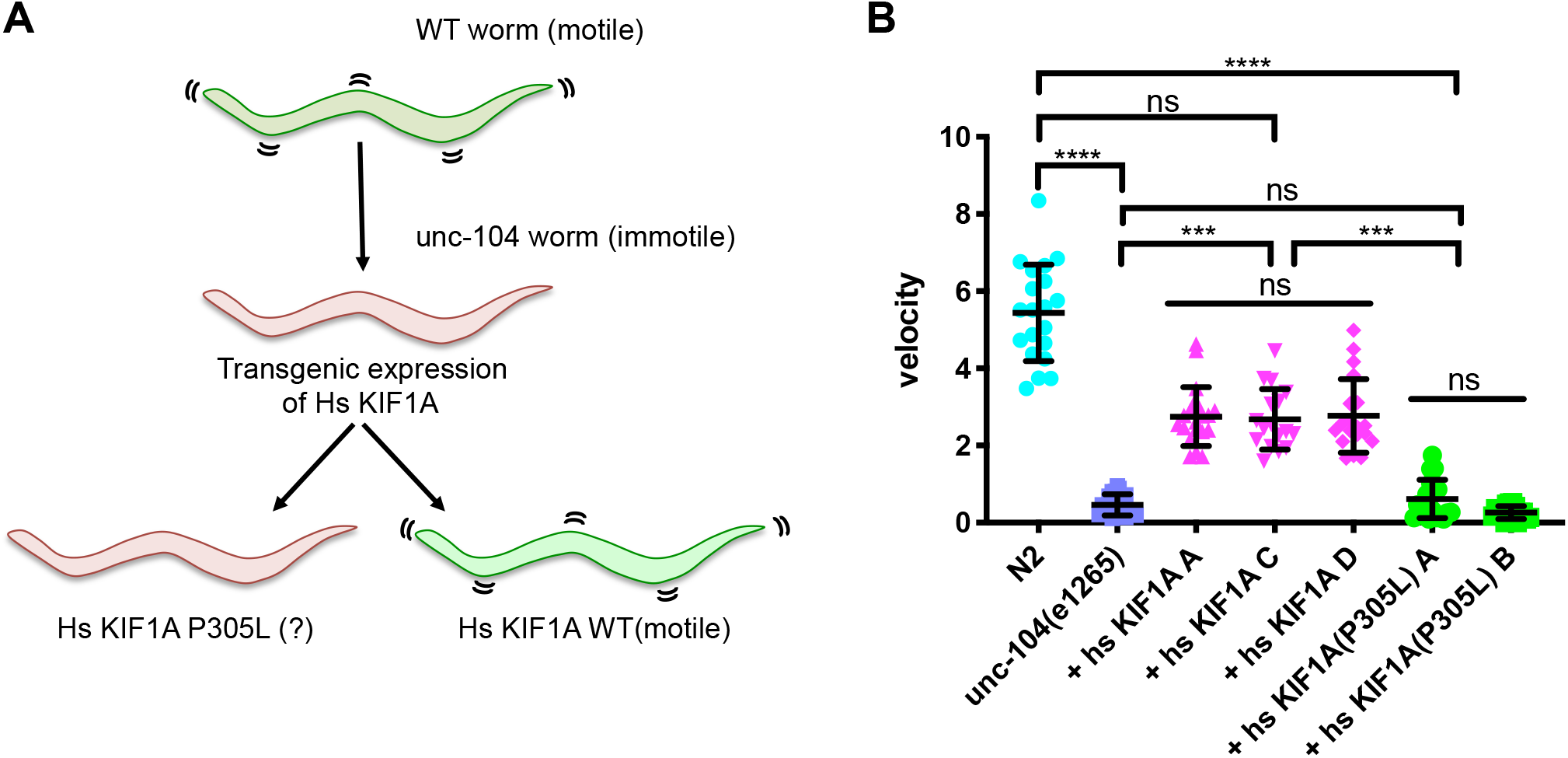
Characterization of KIF1A^P305L^ in a *C*.*elegans* model. **(A)** Cartoon schematic showing the experimental approach. Ablation of the *C*.*elegans* homologue of KIF1A, unc-104, leads to the uncoordinated phenotype in which mutant worms are unable to move robustly. In this genetic background, introduction of a WT human KIF1A gene partially rescues animal movement. Human KIF1A^P305L^ was introduced and assayed for its ability to rescue animal motility. **(B)** The velocity of worm movement is plotted for each genetic background. Three independent lines were assayed for human KIF1A and two independent lines were assayed for human KIF1A^P305L^. Note that while human KIF1A rescues worm movement to approximately half of the normal worm velocity, KIF1A^P305L^ is unable to do so. Data points represent individual worms. Statistical differences between conditions were assessed by Kruskal-Wallis test. Not significant: ns, P ≤ 0.001: ***, P ≤ 0.0001: ****.

**Figure 2.**
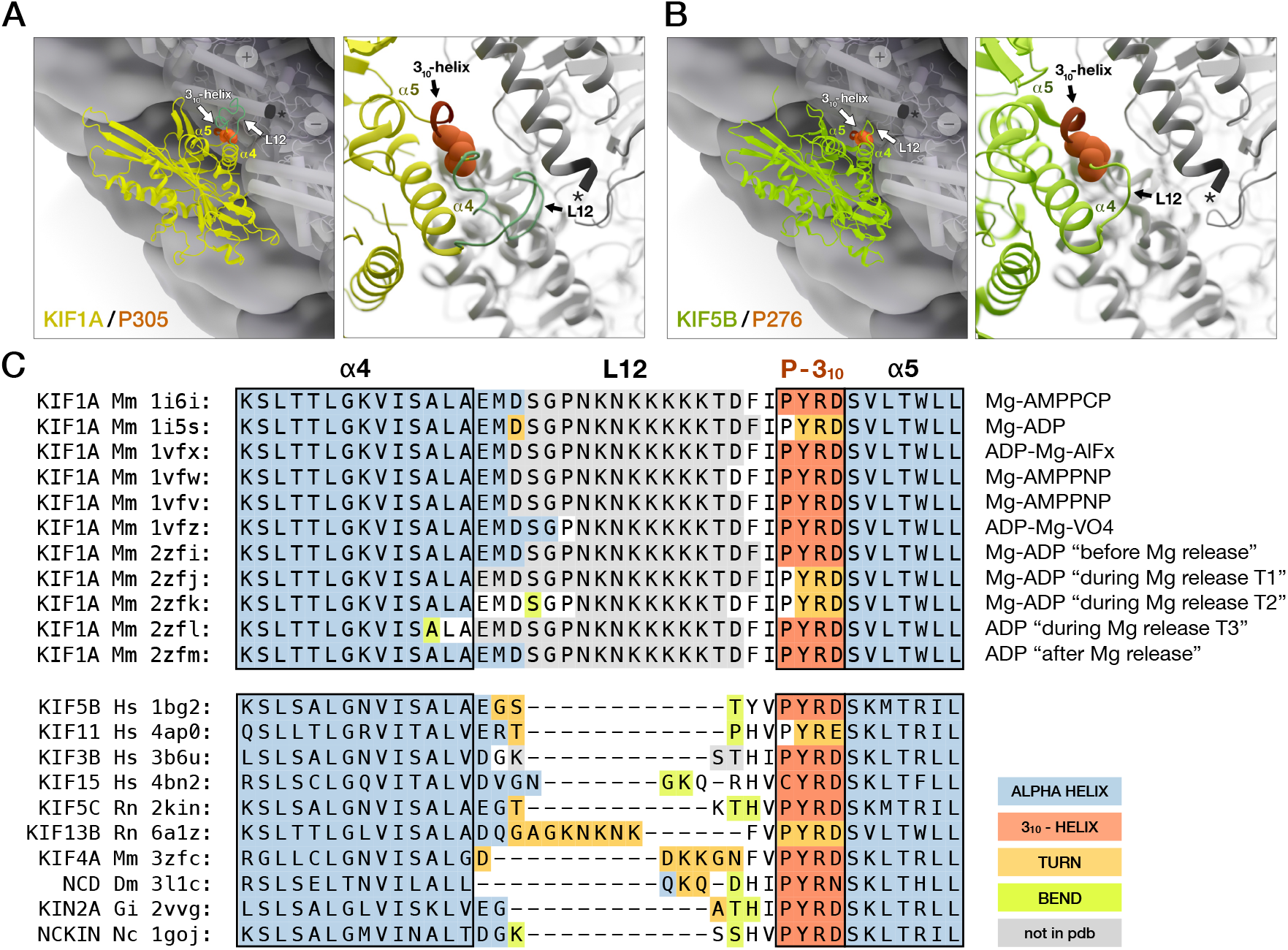
Visualization of the location of P305 residue and 3_10_-helix within the kinesin motor domain. **(A-B)** KIF1A (yellow; pdb: 1vfx), with K-loop/L12 (cyan-green; 288-303) modeled via Swiss-Model (Waterhouse et al., 2018) and shown for comparison to loop L12 in KIF5B (green; pdb: 4hna). */star, beta-tubulin C-terminus. 3_10_-helix (red); surrounding alpha-helices: α4, α5. P305 in KIF1A, and corresponding P276 in KIF5B, are shown as sphere representations (orange). **(C)** Sequences of motor domains with available structures were aligned using ClustalW (Chenna et al., 2003). Residues surrounding the 3_10_-helix are shown. Annotation (left): gene name, species abbreviation, pdb id. (right) KIF1A magnesium (Mg)-nucleotide bound form. Species: *Hs/H*.*sapiens*, Rn/*R*.*norvegicus*, Mm/*M*.*musculus*, Dm/*D*.*melanogaster*, Gi/*G*.*intestinalis*, Nc/*N*.*crassa*. Assignment of secondary structure was done using Procheck (Laskowski et al., 2018; Laskowski et al., 1996) according to the method of (Kabsch and Sander, 1983). Background coloring: alpha-helix (blue), 3_10_-helix (orange), hydrogen-bonded turn (yellow), bend (green), without assignment (white); residues not available in pdb (grey).

Prior mutagenesis studies of charged residues within this motif suggest it plays a critical role in the kinesin-MT interaction, though mutation of the analogous proline in KIF5B (P276) to alanine, which is less bulky than leucine, has little effect on the K_m_ for MTs or on the MT-gliding rate of the motor (23). It is unknown how this KIF5B (P276A) mutant motor behaves under resistive load. In certain Mg-ADP bound KIF1A structures, residues in the PYRD motif deviate from the 3_10_-helix and adopt other non-helical conformations (e.g. hydrogen-bonded turn; Fig. 2C), which we discuss further in Results below. Strikingly, KAND mutations are found in three of the four residues that make up this 3_10_-helix in human KIF1A (P305L, Y306C, R307G/P/Q), and HSP mutations in two of these residues in human KIF5A (P278L, R280H/C/L), and one in KIF1C (R301G) (8, 16, 24-26). These observations highlight the importance of this structural element in kinesin function and human disease.

Here we perform a comprehensive analysis of the P305L variant in KIF1A using genetic, biochemical, and single-molecule methods. We find that the mutation cannot rescue the loss of KIF1A activity in a genetic model system, implying it results in a complete loss of function in living cells. While the mutation reduces the velocity, run-length, and force generation of homozygous mutant motors *in vitro*, these effects are dwarfed by a massive defect in the MT association rate of the motor, particularly in the weak-binding ADP-state of the motor. These results are consistent with a role of the 3_10_-helix in modulating the activity of kinesin’s L12/K-loop, a critical MT-binding element. Interestingly, mutation of the analogous residue in the orthogonal kinesin-1 (KIF5) family, results in an even larger defect in MT-association, explaining the molecular basis of the HSP mutation P278L found in KIF5A. Together, our results lend support to the hypothesis that the conformation of the L12/K-loop, not just its mere presence (27, 28), is critical to enhance the motor’s MT affinity during the motor’s MT-binding cycle. More broadly, we identify a highly conserved 3_10_-helical element, immediately following loop L12 in kinesin, that is critical for the affinity of the motor for MTs. With its effect on MT affinity, it contributes to the motor’s high MT on-rate and its ability to move and generate force, highlighting an understudied aspect of the kinesin-MT interface.

## Results

### A genetic model reveals KIF1A^P305L^ behaves as a null mutation in vivo

To characterize the molecular defects associated with the P305L mutation in KIF1A, we first utilized a well-characterized genetic model system in *C. elegans*. In this system, the worm homologue of KIF1A, *unc-104*, is inactivated by a point mutation within the cargo-binding PH domain located in the distal C-terminal tail region of the motor (*unc-104*(e1265)). The inability of the motor to bind to cargo results in degradation of the protein via the ubiquitin pathway (29). The loss of KIF1A-cargo binding and reduction in motor protein levels results in the ‘uncoordinated’ movement phenotype in which mutant worms are unable to move on agar plates (Fig. 1B, mean velocity for wild-type (WT) and *unc-104*(e1265) 5.4 ± 1.2 and 0.5 ± 0.3 mm/min, respectively). We have previously shown that introduction of the human KIF1A gene into *unc-104(e1265)* background results in restoration of movement to near WT levels (14), indicating that the human gene can largely compensate for unc-104’s cellular functions in worms. Indeed, single-molecule studies with recombinant proteins show that UNC104 and KIF1A have similar motile and force-generation properties (30).

We created three independent rescue lines using WT human KIF1A and two independent rescue lines using human KIF1A^P305L^ and assayed the resulting animals for movement phenotypes on agar plates. Consistent with our previous results, expression of human KIF1A rescued animal movement to approximately 50% of WT velocity (Fig. 1B). While this rescue is weaker than we previously observed (14), the phenotype may correlate with gene expression levels, which are likely variable in our lines. In contrast to WT KIF1A, transgenic expression of KIF1A^P305L^ did not rescue movement defects (Fig. 1B), indicating that the P305L mutation results in near complete loss of KIF1A motor function within living cells. These results strongly suggest that the KAND phenotype in humans is driven by loss of KIF1A motor function in cells.

### The P305L mutation negatively impacts KIF1A’s mechanochemistry

The evolutionarily conserved P305 residue lies immediately C-terminal to the K-loop/L12 within the KIF1A motor domain (Fig. 2). The K-loop/L12 has not been directly visualized in any prior structural studies of KIF1A, suggesting high structural heterogeneity in both MT-bound and unbound states, in contrast to the shorter loop L12 in KIF5B, which has been resolved in published structures (31). We modeled the missing KIF1A K-loop, which is shown for visual comparison in the context of P305, to L12 in KIF5B and its equivalent residue P276 (Fig. 2A, B). In our pseudo model of KIF1A, the K-loop is visualized with its 12 additional residues, 6 of which are positively charged lysines (Fig. 2C); it also bears a flanking proline distal to P305, at position 292, which does not acquire secondary structure in published studies (Fig. 2C) and is not observed to be mutated in KAND patients (Boyle et al., 2020). The additional length and charge of the K-loop may bring it in close proximity to the highly negatively charged (acidic) C-terminal tail of β-tubulin (Fig. 2A, star), and confer motor properties distinct from other kinesins. The P305 residue in KIF1A (Fig. 2A), and P276 in KIF5B (Fig. 2B), as well as equivalent proline residues in other kinesins (not shown), orient similarly with respect to tubulin. This suggests functional conservation across diverse kinesin families, which do not bear lengthy loops in the L12 region. Mutation of P305, or the equivalent proline in other kinesins, likely disrupts the ability of the region to fold as a labile 3_10_-helical structure. In the case of KIF1A, the mutation may disturb favorable orientation of the K-loop with respect to the C-terminal tail of β-tubulin, or it may interfere with unique properties conferred upon the motor by this extended and charged element during its mechanochemical cycle.

We aligned all available high-resolution structures of KIF1A spanning several carefully studied states corresponding to the motor’s nucleotide hydrolysis cycle (Fig. 2C). The 3_10_-helix is present in most states observed. Notably, in two KIF1A-Mg-ADP transition states, corresponding to the motor undergoing Mg-release upon binding to the MT (2zfj/T1 and 2zjk/T2; Fig. 2C; (Nitta et al., 2008)), the proline-phi torsion angle for P305 is three standard deviations outside normal (not shown). Proline’s atypical restrictions in phi-psi space arise from its 5-membered ring, which limits rotation about the N-αC bond in the polypeptide backbone. Thus, what corresponds to a captured state with PYRD in non-helical form and an unusually high proline-phi torsion angle, may be indicative of a structural element under strain, or allosteric regulation. Altogether, this would suggest the unique PYRD 3_10_-helix may act as a labile structural element, or temporal regulatory switch, that is critical for specific steps which may couple to the motor’s mechanochemical cycle, or for helping to tether the motor head when the kinesin is under load. The PYRD/E motif also adopts a 3_10_-helical conformation in structures from other kinesin families and species examined (Fig. 2C).

Based on this analysis, it is possible that mutation of P305 could directly impact KIF1A’s interaction with the MT, although pseudo-atomic models fit to electron density of KIF1A bound to MT would not suggest direct contact (Atherton et al., 2014). We have previously reported that some human disease mutations within the KIF1A motor domain do not disrupt the motor’s mechanochemical cycle, but rather impair the autoinhibition mechanism of the full-length motor (5, 32, 33) leading to hyperactivation of motor activity, which also appears to lead to KAND (14). Thus, we hypothesized that the P305L mutation would either directly impact the motor’s mechanochemical cycle, or possibly, disrupt the autoinhibition mechanism of KIF1A. Importantly, these hypotheses have disparate predictions; the first would result in decreased motor activity, while the latter increased motor activity. Another possibility we considered is that the mutation disrupts the three-dimensional folding of the motor domain, leading to unstable or aggregated protein, and thus a loss of KIF1A activity, or even possibly a toxic gain-of-function, in cells.

To distinguish between these possibilities, we first turned to single-molecule assays utilizing full-length, recombinant, purified KIF1A motors (14). The full-length motor has a relatively low MT-landing rate, compared to the more widely studied tail-truncated and constitutively active motor. This is presumably due to autoinhibition of the full-length motor, although mechanistic details of KIF1A’s mechanism of autoinhibition remain unclear (5, 32, 33). Using baculovirus expression, we purified full-length KIF1A and KIF1A^P305L^ and assessed their structural integrity using size-exclusion chromatography (SEC). Both purified motors eluted from the SEC column indistinguishably (Fig. 3A). These results argue against a deleterious effect of the P305L mutation on protein folding or stability. Next, we utilized multi-color, single-molecule total internal reflection microscopy (TIRF-M) to directly visualize the motor activity of our purified motors. As we previously observed, WT KIF1A moved long distances along MTs in the presence of ATP (Fig. 3B). Surprisingly, we also observed long-distance movement of KIF1A^P305L^ (Fig. 3B), ruling out the possibility that the mutation abolishes motor activity. We measured the velocity of these movements and observed that the P305L mutation lead to an ∼25% decrease in velocity (1175 ± 528 vs. 876 ± 479 nm/s [±SD] for KIF1A and KIF1A^P305L^ respectively, Fig. 3C). In addition, we noted an ∼50% decrease in motor run-lengths (2.5 ± 0.3 vs. 1.2 ± 0.2 µm for KIF1A and KIF1A^P305L^ [±SD], respectively, Fig. 3D). These data reveal that the P305L mutation does not disrupt protein folding or stability, but rather results in impaired motility of the KIF1A motor.

**Figure 3.**
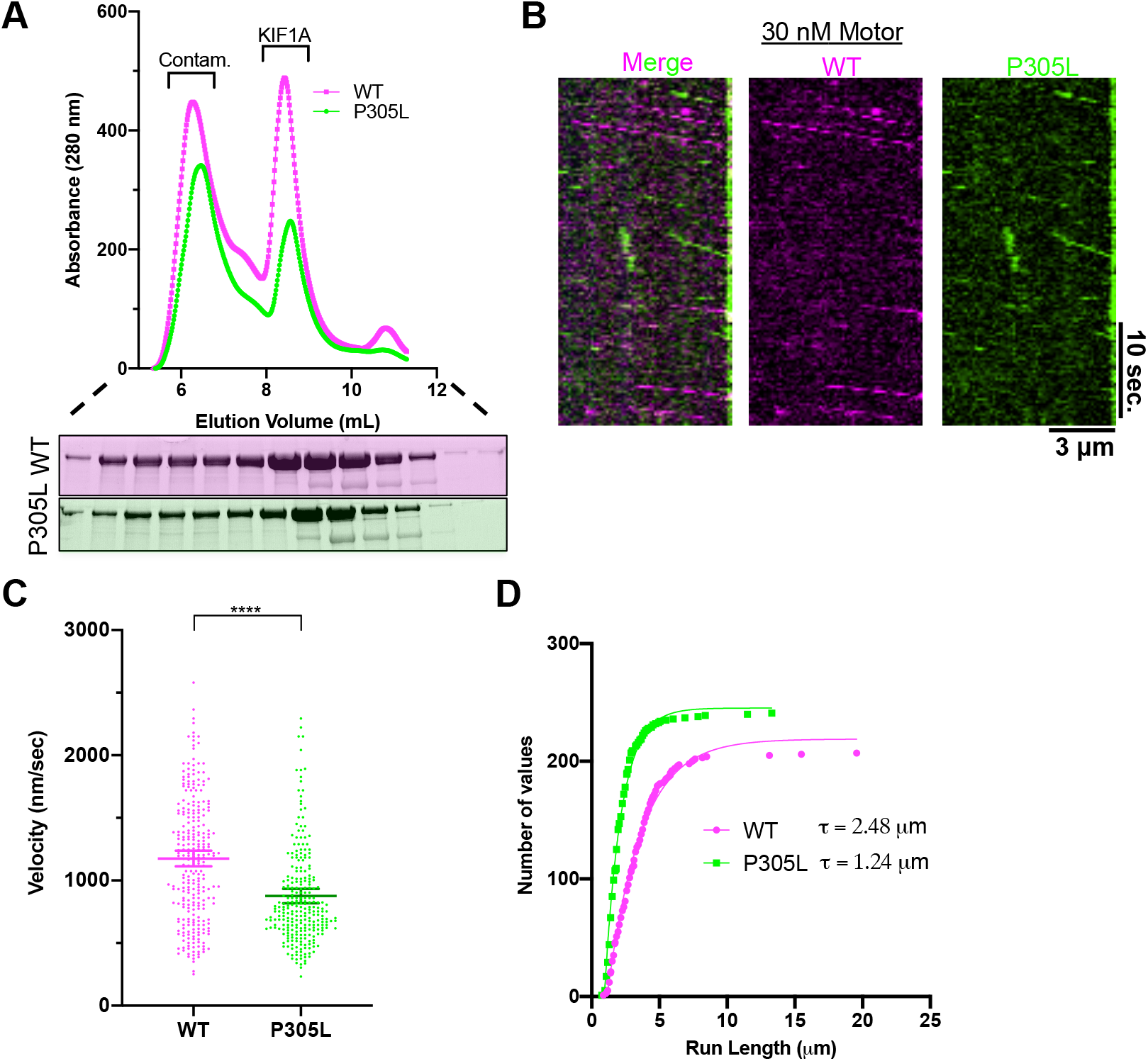
Characterization of the KIF1A^P305L^mutation using full-length motors. **(A)** SEC chromatogram showing elution profiles of full length KIF1A (magenta) and KIF1A^P305L^(green). Elution positions of prep contaminants (Contam.) and full-length KIF1A are noted. Below: Coomassie stained SDS-PAGE gel of the column elution fractions. **(B)** Kymograph from TIRF-M movie showing both KIF1A (magenta) and KIF1A^P305L^ (green) motors moving along the same MT. Equal concentration of each motor was added to the TIRF chamber. **(C)** Measured velocity distribution of full-length KIF1A motors (N = 2 independent trials and n = 280, 270 motors from KIF1A and KIF1A^P305L^ respectively). **(D)** Cumulative distribution plot of the measured run-lengths of WT and P305L motors. Connecting lines show fit to a one-phase exponential decay function (R^2^= 0.99, 0.98 for KIF1A and KIF1A^P305L^ respectively). The characteristic run-length derived from the fits (τ) is displayed (N = two independent trials, n = 207, 241 for KIF1A and KIF1A^P305L^ respectively).

### The P305L mutation most dramatically impacts KIF1A’s initial MT interaction

While prior studies have found defects in the velocity and run-lengths of mutant motor proteins *in vitro* (34, 35), it is currently unclear how important these biophysical parameters actually are for motor protein function *in vivo*. For instance, while KIF1A is ultra-processive on bare MTs *in vitro*, recent evidence suggests that non-motor MAP proteins (36), as well as other motor proteins, on the same MT (37), can dramatically affect the motile parameters of KIF1A and other motor proteins. Given the relatively modest effects we observed for the P305L mutation on the velocity and run-lengths of full-length KIF1A motors *in vitro* (Fig. 3), we wondered whether these defects could account for the dramatic phenotypes observed in humans with this mutation. The aforementioned results are complicated by the fact that we still do not fully understand how full-length KIF1A motors are autoinhibited, or how that autoinhibition is relieved *in vivo* or *in vitro*. Thus, we turned to a well-characterized KIF1A construct in which the entire tail domain is removed, and the motor is stably dimerized via fusion to a leucine-zipper motif (KIF1A 1-393-LZ). This KIF1A construct is constitutively active and has a high MT landing rate due to the lack of autoinhibition elements located in the tail domain (5, 32, 33).

We purified KIF1A 1-393-LZ proteins from baculovirus infected insect cells and assessed protein folding via SEC. Both WT and P305L motors eluted similarly from the column (Fig. 4A), further indicating that the P305L mutation does not grossly perturb the folding of the KIF1A motor domain. We assessed the ability of these motors to interact with, and translocate along MTs using our multi-color TIRF-M assay. We mixed differentially labeled WT and P305L mutant motors and observed their movement within the same TIRF chamber, along the same MTs, providing a robust internal control for the assay conditions. Because the MT-binding rate of this truncated motor is very high due to removal of the autoinhibition mechanism, we utilized an ∼30-fold lower concentration of WT motor as compared to our assays with full-length KIF1A (Fig. 3). However, at these low concentrations, while we could observe many motile WT motors, we could not detect any mutant motors moving along MTs (not shown). Therefore, we increased the concentration of KIF1A^P305L^ motors by 10-fold (as compared to WT, Fig. 4B). In these conditions, we could observe the movement of both types of differentially-labeled motors moving along the same MTs in the chamber, although the number of moving KIF1A^P305L^ motors was clearly lower than that of the WT motors (Fig. 4B).

**Figure 4.**
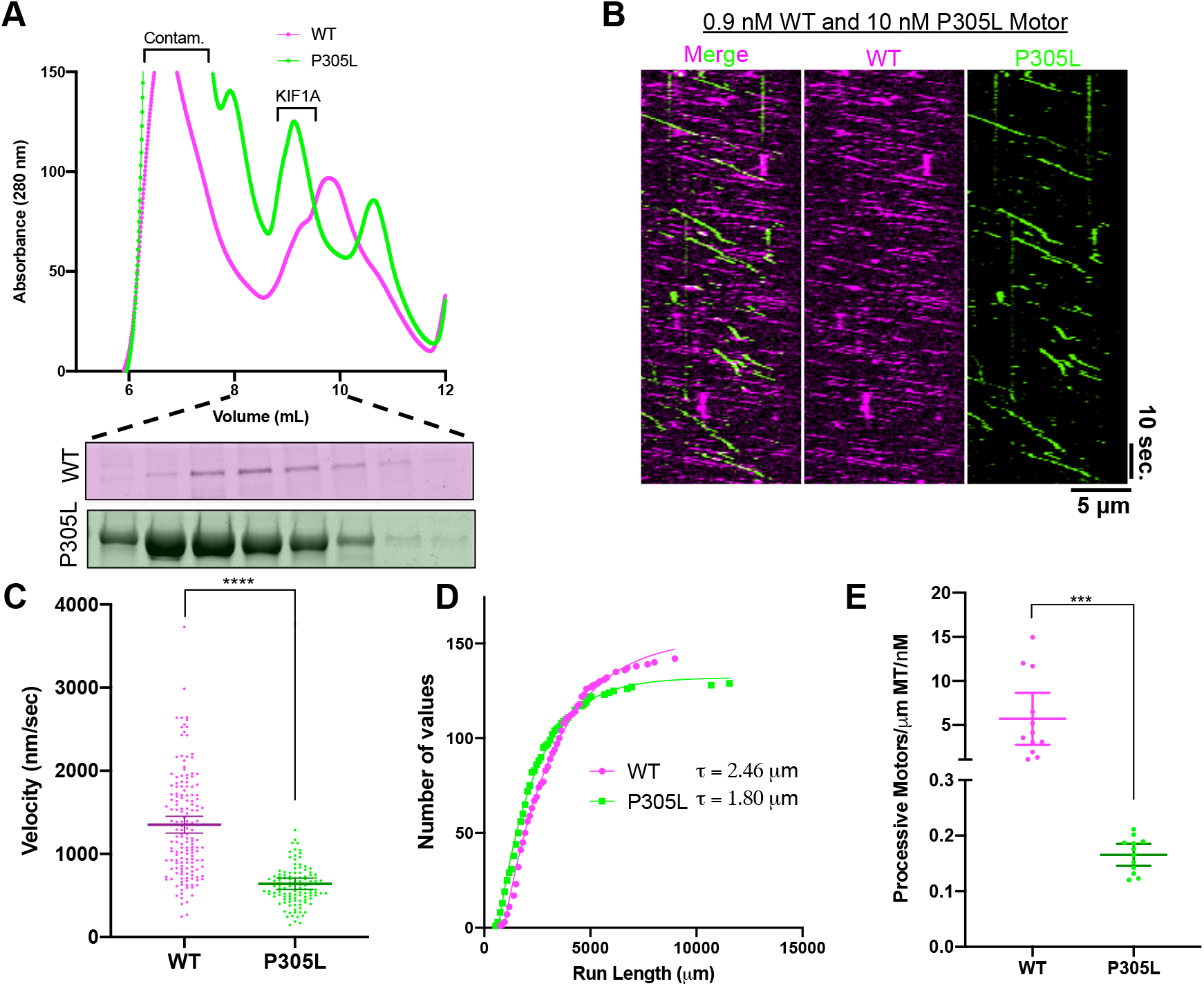
Characterization of the KIF1A^P305L^ mutation using truncated, constitutively active motors. SEC chromatograms for WT (magenta) and P305L truncated (green), dimerized motors (1-393-LZ). Elution positions of prep contaminants (Contam.) and truncated KIF1A are noted. Below: Coomassie stained SDS-PAGE gel of the column elution fractions encompassing the motor peak. **(B)** Kymograph from a TIRF-M movie showing both KIF1A (magenta) and KIF1A^P305L^(green) moving along the same MT. Note the difference in motor concentration. **(C)** Measured velocity distribution of 1-393-LZ KIF1A motors (N = 2 independent trials and n = 161, 116 motors from KIF1A and KIF1A^P305L^ respectively). **(D)** Cumulative distribution plot of the measured run-lengths of WT and P305L 1-393-LZ motors. Connecting lines show fit to a one-phase exponential decay function (R^2^= 0.99 for both KIF1A and KIF1A^P305L^ respectively). The characteristic run-length derived from the fits (τ) is displayed (N = two independent trials, n = 142, 129 for KIF1A and KIF1A^P305L^ respectively). **(E)** Measured number of processive motors per unit length of MT and time (N = two independent trials, n = 12 MTs analyzed each for KIF1A and KIF1A^P305L^). For all panels, P ≤ 0.001: ***, P ≤ 0.0001: ****.

Inspection of the resulting kymographs revealed that the slopes of motile KIF1A^P305L^ appeared more shallow, indicating the mutant motors were moving at reduced velocities, consistent with our results with full-length motors (Fig. 3). Quantification of motor velocities confirmed that the P305L mutation impaired motor velocity by approximately 50% (1352 ± 650 vs. 641 ± 368 nm/s [± SD] for KIF1A and KIF1A^P305L^, respectively, Fig. 4C). These results are consistent with the results we obtained with full-length motors (Fig. 3). Additionally, we measured the run-lengths of the truncated motors and observed a more modest ∼25% decrease for the mutant motors (2460 ± 67 vs. 1803 ± 210 nm [±SD] for KIF1A and KIF1A^P305L^ respectively, Fig. 4D), similar to our results with full-length KIF1A motors (Fig. 3). It is interesting to note that the magnitude (but not the trend) of the effects of the P305L mutation on velocity and processivity differ between full-length (Fig. 3) and tail-truncated motors (Fig. 4). We hypothesize that these biophysical differences are attributable to biochemical distinctions bestowed by the autoinhibition mechanisms of full-length motors.

The initial motor-MT interaction of the tail-truncated motor is not complicated by the autoinhibition mechanism present in full-length motors, allowing us to directly compare the number of motile motors on individual MTs within the same imaging chamber. As inferred by our initial observations of differential motor concentrations required for observation of motile molecules (see above), we measured a very dramatic decrease in the numbers of moving molecules along MTs for the P305L construct (Fig. 4E). This ∼97% (71-fold) drop in motor landing rate was much larger than the ∼50% decrease in velocity or ∼25% drop in processivity. Thus, the P305L mutation results in relatively modest defects in KIF1A’s motor biophysical output once it is moving processively along MTs, but a precipitous decline in the initial motor-MT interaction, implying that the KAND phenotype (as well as the null phenotype we observed in our worm model) may largely be due to the inability of the mutant motor to get onto the MT. Interestingly, the K-loop (L12) of KIF1A has been reported to be critical for the motor’s MT on-rate, but not for its movement (3), suggesting the largest observed defect of KIF1A^P305L^ could be the result impaired K-loop function.

### The P305L mutation impairs the ability of the K-loop to enhance the motor-MT interaction during the ADP state

It has been previously reported that the K-loop is critically important for the KIF1A-MT interaction during the weak binding state of the motor’s mechanochemical cycle (the ADP-bound state) (3, 28, 38). Swapping the KIF1A K-loop with the less positively charged L12 from KIF5C increased the dissociation constant (K_d_) for MTs by ∼4-fold (28), while single-molecule studies confirmed that mutation of the K-loop abolished the interaction of ADP-bound KIF1A with MTs (3). The interpretation of these results is that the positively charged K-loop interacts with the nearby negatively charged C-terminus of β-tubulin to facilitate the KIF1A-MT interaction (Fig. 2A). Thus, we examined the effect of the P305L mutation on the interaction of truncated KIF1A dimers, in their weak binding ADP-state, with MTs using single molecule imaging. We again noted a large difference in the concentration-dependence of motors interacting with MTs between WT and KIF1A^P305L^ motors (Fig. 5A). Quantification of the motor landing rates revealed an ∼ 83% reduction in landing rate for KIF1A^P305L^ motors in the ADP-state (Fig. 5B). Measurement of the motor dwell-times on MTs in this state revealed a much more mild ∼ 27% decrease in dwell-time for KIF1A^P305L^ versus WT motors in ADP (Fig. 5C). We also measured the steady-state ATPase rate of the motors and observed an ∼250-fold increase in the K_m_ for KIF1A^P305L^ (Fig. 5D), further demonstrating a strong defect in MT-association. Since the K_m_ approximates the MT affinity of kinesin in the ADP state (23), we conclude that the P305L mutation primarily impacts the motor’s MT association rate in the ADP state.

**Figure 5.**
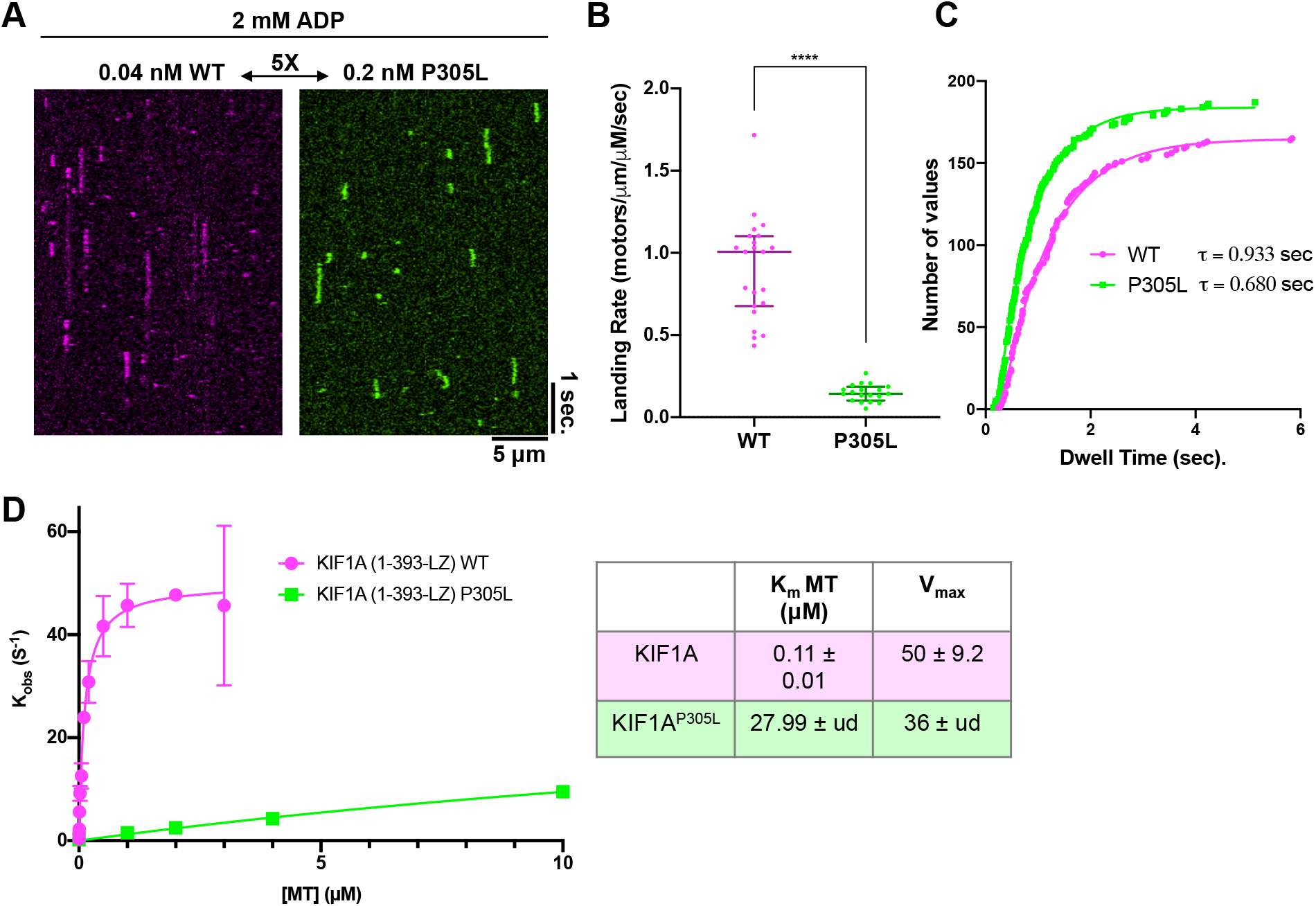
Characterization of the weak MT-binding state of KIF1A. **(A)** Kymographs from TIRF-M movies showing binding and dwell events of 1-393-LZ motor constructs in 2mM ADP. Note the different protein concentrations used. **(B)** Quantification of the landing rate for KIF1A and KIF1A^P305L^motors (N = two independent trials, n = 12 MTs analyzed each for KIF1A and KIF1A^P305L^). **(C)** Cumulative distribution plot of the measured dwell-times of WT and P305L 1-393-LZ motors. Connecting lines show fit to a one-phase exponential decay function (R^2^= 0.99 for both KIF1A and KIF1A^P305L^ respectively). The characteristic dwell-time derived from the fits (τ) is displayed (N = two independent trials, n = 165, 187 for KIF1A and KIF1A^P305L^ respectively). **(D)** Steady-state ATPase assays for KIF1A and KIF1A^P305L^. Plotted are the mean values from at least 2-3 independent trials per concentration of MTs, error bars are S.E.M. Non-linear fits to the Michaelis-Menten equation are shown, and the calculated Km and Vmax are displayed in the table to the right with 95% CI errors shown. Note the errors of the fit for KIF1A^P305L^ cannot be accurately calculated by this method (ud: undetermined).

These results are consistent with a prior model that proposes that the K-loop primarily functions to endow KIF1A with a high MT association rate during its weak binding (ADP) state (3). However, our results reveal that the presence of the positively charged region of the K-loop is not sufficient for its function, and we posit that the conformation of loop L12, transiently constrained via an adjacent labile 3_10_-helix and modulated by the atypical properties of the proline residue, at the N-terminal cap of the 3_10_-helix, is required for this function. Strikingly, three of the four residues that make up the PYRD 3_10_-helix are mutated in KAND patients (8), and we hypothesize that such mutations (Y306C, R307G/P/Q), which may destabilize this unique helix, are likely to exhibit effects on the KIF1A-MT interaction that are similar to P305L. However, patients with mutations in different positions in the motif, and multiple patients with mutations in the same position, exhibit varying degrees of disease severity (8), which suggests nuanced effects on the motor’s MT functions, or developmental or other effects *in vivo*.

### The functional role of the L12-proximal 3_10_-helix is conserved in different classes of kinesin motors

The P305 residue in KIF1A is part of a highly conserved PYRD/E motif found in many other classes of kinesin motors in humans and other organisms (Fig. 2). This motif adopts a 3_10_-helical conformation in most of the published kinesin structures examined. In some structures, residues in the motif deviate and adopt other non-helical conformations (Fig 2C). The immediate proximity to loop L12, the high sequence conservation, and the preference to adopt a 3_10_-helical conformation, implies that this element plays an important role in the basic kinesin translocation mechanism. The canonical kinesin-1 family (KIF5A, KIF5B, KIF5C) is probably the most well studied kinesin motor, and members of this family also contain the absolutely conserved PYRD motif directly C-terminal to L12 within the motor domain (Fig. 2C). Notably, L12 in the KIF5 family does not contain a highly charged stretch of lysine residues like KIF1A, resulting in a shorter L12 (Fig. 2). We mutated the analogous residue in KIF5B, P276, to leucine to assess the effects of this mutation on KIF5B’s motile properties. We measured the binding of a tail-truncated, fluorescently tagged KIF5B motor (amino acids 1-420) to MTs in the presence of ATP. The WT motor bound robustly to MTs in the presence of ATP (Fig. 6A) and moved processively (Fig. 6B), as previously characterized by many labs. Strikingly, KIF5B^P276L^ showed little observable binding to, or movement along, MTs at two different concentrations spanning an order of magnitude (Fig. 6A, B). Indeed, quantification of the fluorescence intensity of kinesin on MTs in ATP revealed an enormous decrease of ∼4-5 orders of magnitude in fluorescence intensity for the KIF5B^P276L^ motor (Fig. 6C), to near background levels, indicating an almost complete loss of the kinesin-MT interaction. The near complete lack of binding to MTs was distinctly worse than the effects we observed with KIF1A^P305L^ (Figs. 3-5), prompting us to investigate if KIF5B^P276L^ was even capable of interacting with MTs. However, in the presence of the non-hydrolyzable ATP analogue, AMP-PNP, we observed robust binding of KIF5B^P276L^ to MTs (Fig. 6A), indicating that the motor is still capable of interacting with MTs in this particular state. Together, our data reveal that the conserved proline adjacent to L12 in multiple kinesin families plays a crucial role in L12’s ability to facilitate the interaction of the kinesin motor domain with the MT surface, particularly in the ADP state of the mechanochemical cycle.

**Figure 6.**
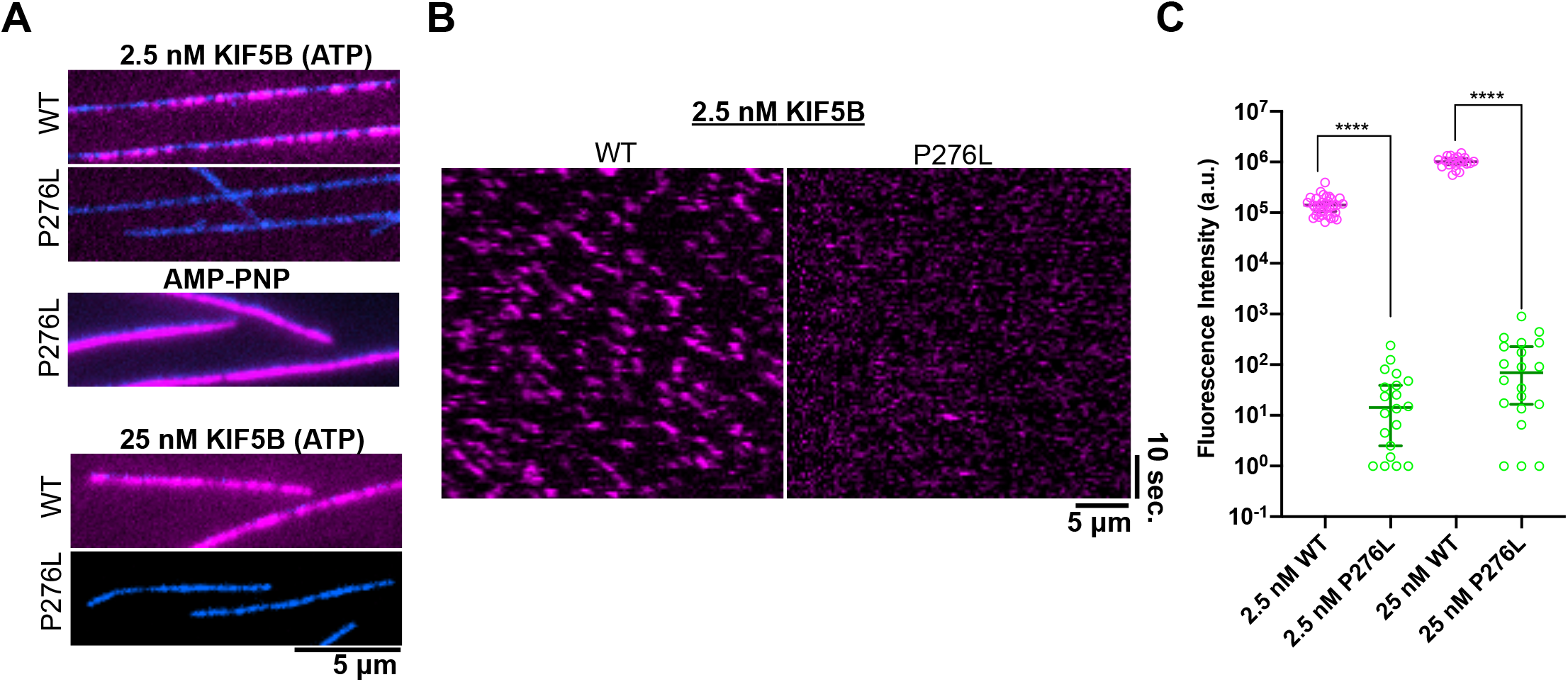
Characterization of conserved proline mutation in KIF5. **(A)** Identically scaled TIRF-M images showing MTs (blue) and KIF5 molecules (magenta) in ATP at two different protein concentrations. Note the lack of MT binding for KIF5^P276L^. KIF5B^P276L^ binds robustly to MTs in the presence of AMP-PNP, demonstrating the motor is still proficient at binding. **(B)** Kymographs from TIRF-M movies of KIF5B motors in ATP demonstrate processive movement along MTs (diagonal lines) for KIF5B, but not for KIF5B^P276L^. Only transient binding events are observed for KIF5B^P276L^. **(C)** Quantification of mean fluorescence intensity of KIF5B motors along MTs in the presence of ATP at two different protein concentrations (N = two independent trials, n = 20-35 MTs quantified per condition.

### The P305L mutation impairs KIF1A’s ability to generate force along MTs

After determining the effects of the P305L mutation on the KIF1A-MT interaction and KIF1A motility in the absence of load, we wondered if the mutation might have any effects on the ability of KIF1A to generate force along MTs, which is of particular importance for the KIF1A-powered displacement vesicular cargo within living cells. To assess any effects on force generation, we performed single-molecule optical tweezers assays (7, 39, 40) with our human KIF1A proteins. In a parallel study (30), we have shown that full-length rat KIF1A and tail-truncated rat KIF1A (1-393)-LZ both detach at an average load of ∼2.7 pN and then rapidly reattach to the MT to resume motion, resulting in a sawtooth-like force-generation pattern in which force-generation events cluster tightly together (30). We find that human KIF1A (1-393-LZ), which differs from a similar rat KIF1A construct by a single amino acid (valine instead of an isoleucine at residue 359), also exhibits a sawtooth-like force-generation pattern (Fig. 7A), but detaches at a reduced average force of 2.2 [1.9, 2.5] pN (median [quartiles], Fig. 7E). In contrast to human WT KIF1A, KIF1A^P305L^ detaches at a ∼4-fold reduced force of [0.5, 0.8] pN (Fig. 7B, E), revealing an even greater effect on force generation than on velocity and processivity (Figs. 4 and 7). These results are consistent with our observations that the mutation dramatically impinges on the motor’s ability to bind to the MT (Figs. 3 and 4). Thus, we conclude that the 3_10_-helix plays a critically important role in KIF1A’s ability to generate force and move under resistive load. Considering that L12 is still intact in the mutant motors, the data further suggest that the conformation of L12, not its mere presence, is crucial for its role in promoting the kinesin-MT interaction during force generation.

**Figure 7.**
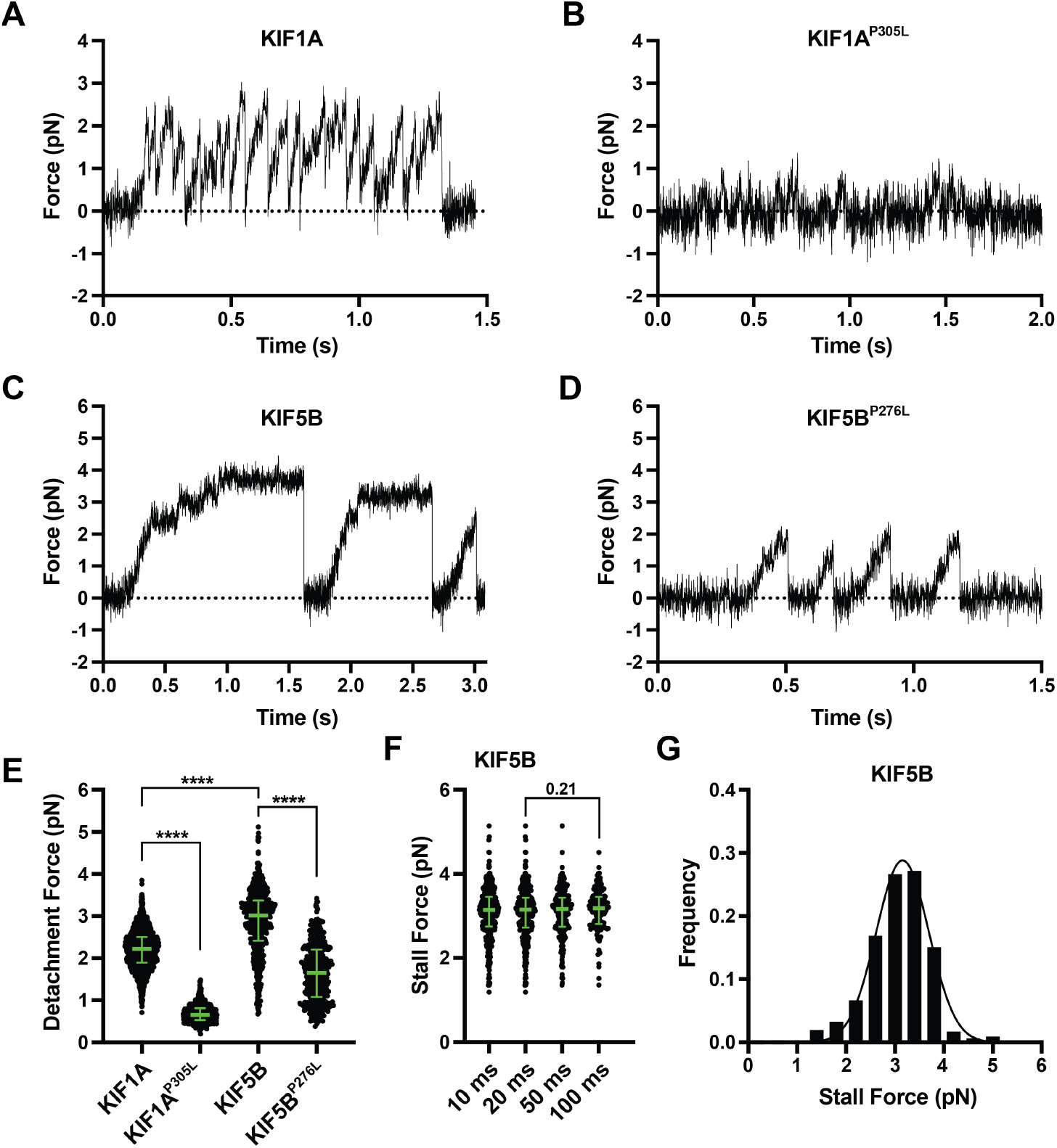
Characterization of the force-generation properties of KIF1A^P305L^ and KIF5B^P276L^. **(A-D)** Representative force vs. time records of bead movement driven by single molecules of **(A)** WT KIF1A(1-393)-LZ, **(B)** KIF1A^P305L^(1-393)-LZ, **(C)** WT KIF5B(1-420), and **(D)** KIF5B^P276^(1-420). **(E)** Detachment forces. Green bars indicate the median values with quartiles. WT KIF1A(1-393)-LZ: 2.2 [1.9, 2.5] pN, *N* = 1737; KIF1A^P305L^(1-393)-LZ: 0.7 [0.5, 0.8] pN, *N* = 637; WT KIF5B(1-420): 3.0 [2.4, 3.4] pN, *N* = 603; KIF5B^P276^(1-420): 1.7 [1.1, 2.2] pN, *N* = 415. **(F)** Maximal force (“stall force”) sustained during a single run against load for a minimum time duration of 10 ms (3.1 [2.7, 3.5] pN, *N* = 405), 20 ms (3.1 [2.7, 3.4] pN, *N* = 381), 50 ms (3.2 [2.7, 3.4] pN, *N* = 262) and 100 ms (3.2 [2.8, 3.5] pN, *N* = 154), respectively. **(G)** Stall force of WT KIF5B(1-420) (≥20 ms criterion).

Finally, to define whether the 3_10_-helix adjacent to L12 is also involved in the force generation of kinesin-1, we performed optical trapping experiments on WT and P276L tail-truncated KIF5B. We find that WT KIF5B shows stalling events as expected for a kinesin-1 motor, but does so at an average force of 3.2 ± 0.5 pN (mean ± SD; ≥20 ms criterion, Fig. 7C, E-G), which is less than the previously reported maximal force generation of 5-6 pN of full-length kinesin-1 and the widely-studied KIF5B (1-560) construct (41-45). Even if we apply a stalling criterion of 100 ms, the stall force remains at 3.2 ± 0.4 pN (Fig. 7F). We hypothesize this difference could be due to the shorter KIF5B (1-420) construct used in this work, but further studies are necessary to determine the exact reason for the lower stall force of this construct. We note that most prior studies on kinesin force generation used non-physiological acidic pH buffers (typically pH ∼6.8), and previous data revealed that the interaction of kinesin with MTs is highly sensitive to pH (46). Thus, the lower stall forces we observe may in part be a consequence of the higher pH buffer (pH 7.2) used in our assays.

Nonetheless, a comparison of the average detachment forces of human WT KIF5B and KIF5B^P276L^ (3.0 [2.4, 3.4] vs. 1.6 [1.1, 2.2] pN, median [quartiles], Fig. 7E) revealed an ∼50% reduction in force generation, demonstrating that the 3_10_-helix segment is also critically important for the force generation of kinesin-1 Our results thus reveal that the highly conserved 3_10_-helical element C-terminal to L12 plays important roles under load in kinesin motors of distant evolutionary lineage. Further, these results provide unifying insight into the molecular mechanism underlying PYRD motif mutations that cause KAND in KIF1A and HSP in KIF5A.

## Discussion

Disruption of long-distance transport by kinesin motor proteins is becoming recognized as a prominent driver of neurological diseases such as HSP, KAND, and Rett syndrome. The sheer number of reported KIF1A mutations resulting in KAND and HSP phenotypes complicates the planning and development of effective therapeutics to treat these disorders. Having a molecular understanding of the defects caused by particular mutations, or clusters of mutations that fall within similar structural regions of the motor would greatly enhance therapeutic strategies. Here we use single-molecule methods to dissect the specific defects in an understudied pathological variant of KIF1A, P305L, demonstrating that single-molecule observation, along with traditional bulk biochemical assays and genetic rescue experiments, is a powerful platform to understand the impacts of disease mutations on molecular function. Our work surprisingly revealed that a highly conserved element (adjacent to loop L12 in the kinesin motor core) that is absolutely critical for the kinesin-MT interaction, harbors a labile 3_10_-helical conformation and is a structural locus susceptible to recurring human disease mutations in kinesins.

### Functional conformation of kinesin’s L12/K-loop may be critically mediated by the adjacent 3_10_-helix

Using deletion or replacement mutagenesis, several studies have found that the presence of the highly charged K-loop within the KIF1A motor domain is critical for its function as a long distance transport motor by mediating a high MT-binding rate during the weak-binding (ADP) state of the motor’s hydrolysis cycle (3, 6, 28, 38). However, despite a wealth of structural knowledge of the KIF1A motor domain, the K-loop has never been visualized, presumably due to conformational flexibility. The K-loop follows the main MT-binding element made up of loop 11 (L11) and the α4 helix on the bottom of the kinesin motor domain (28, 47). Crystallography of monomeric KIF1A motor domains in the absence of tubulin (28) found the conformation of α4 changes during the hydrolysis cycle of the motor with the C-terminal end of α4, near the K-loop, elongating in the ADP-Pi state, presumably altering the conformation of the attached K-loop. Release of phosphate results in partial melting of the C-terminal end of α4, again presumably altering the conformation of the attached K-loop to allow for its interaction with the C-terminus of β-tubulin (28). Further, proteolytic mapping of the kinesin motor domain in different nucleotide states also revealed nucleotide-dependent changes in L12, confirming conformational plasticity in this region (27). These data support a model whereby the conformation of the K-loop is dynamic, and directly connected to the ATP hydrolysis cycle of kinesin.

More recent ∼7Å cryo-EM structures of KIF1A bound to MTs, in several states corresponding to the hydrolysis cycle, reveal neither the conformation of the positively charged portion of the K-loop nor the negatively charged C-terminal tubulin tail (47), indicating structural heterogeneity in these regions. However, based on the fitting and calculation of the pseudo-atomic models, these structures did reveal electron density between helix H12 of β-tubulin and parts of the calculated location of the PYRD 3_10_-helical element within KIF1A. While the exact composition of this density in distinct nucleotide states is unclear at present, we hypothesize that it is crucial for mediating a functional conformation of the adjacent K-loop during the weak-binding state of the motor. Interestingly, density between β-tubulin and parts of the PYRD 3_10_-helical element is maintained in all nucleotide states observed, suggesting it may serve as a crucial anchor point to the MT. We hypothesize that this contact may also be critical to orient the conformation of the K-loop and mediate its effects during the landing of KIF1A onto the MT.. There is less clear density connecting the kinesin-1 motor, KIF5A (which was modeled based on KIF5B), to the MT in this same region. Yet, our results reveal that the same proline that initiates the 3_10_-helix segment is also critical for the MT interaction and force generation of KIF5B (Figs. 6-7), and we suggest it likely plays a similar role in mediating L12’s interaction with tubulin in KIF5-family motors and other kinesins (Fig. 2B,C).

An interesting aspect of our data is that the largest effect of the P305L mutation is on the landing rate of kinesin onto the MT, highly similar to deletion or mutation of the K-loop (3, 6). Once mutant KIF1A motors get onto the MT, they are capable of taking many hundreds of steps without dissociating, suggesting the function of the 3_10_-helix may not be as crucial during motor stepping in the absence of resistive load. This is consistent with prior results, which found that mutation of the K-loop strongly perturbs the landing rate, but not processivity of KIF1A (3). However, while we observe more mild defects in velocity and processivity, we see more severe effects on KIF1A force generation under load, suggesting that the mutation also impairs important aspects of the mechanochemical cycle that are sensitive to resistive force. Our data support the notion that the conformation of the K-loop is as important for facilitating the landing of KIF1A onto MTs, as previously reported (3). Based on our data, we favor the interpretation in which altering of the ability of the PYRD/E motif to act as a labile regulatory 3_10_-helix or switch under load, affects allosteric communication between loop L12, nucleotide binding site and MT-binding sites within the kinesin motor. High-resolution structural data of mutant motors would help to further develop this hypothesis.

### Prospects for human disease

Mutations in KIF1A result in a broad range of KAND phenotypes with varying severity (8), suggesting differing disease mechanisms based on the type of mutation in the motor. While some mutations result in a complete loss of motor activity or over-activation of motor activity (12-14), we demonstrate here that the P305L mutation results in a severe loss of MT-binding affinity, but that surprisingly, once mutant motors are properly loaded onto the MT, they are capable of processive movement under unloaded conditions. However, when subjected to load, the P305L mutation severely affects the force output of the motor. All of these results suggest that potential treatment options for KAND may need to be tailored to the specific effects of the particular mutation on the motor’s biophysical outputs. In the case of P305L, therapeutic options that increase the mutant motor’s ability to interact with MTs could be a viable goal. Interestingly, the non-motor MT associated protein MAP9 was recently shown to enhance the landing rate of KIF1A in vitro (36), suggesting that orthogonal endogenous molecules could be useful targets for potential therapeutic interventions.

Multiple residues that make up the short 3_10_-helix are mutated in KIF1A (P305L, Y306C, R307G/P/Q), the related KIF1 family member KIF1C (R301G), and the orthogonal kinesin family member KIF5A (P278L, R280H/C/L), leading to neurodegenerative disease phenotypes in humans (8, 16, 24-26). These findings highlight the critical function of this understudied element of the kinesin motor core. The observation of the same and recurring missense mutation (proline to leucine) in analogous residues of KIF5A and KIF1A (with degrees of disease severity in patients), highlights the importance of this particular residue for the general kinesin motor mechanism. Further, it may suggest a ‘surrogate mechanism’ underlying the specific elasticity of the leucine substitution. It also indicates that therapeutics that address this defect in the kinesin-MT interaction could be taylored to augment or modulate the substitution, and be applicable to several human diseases.

In summary, the single-molecule studies of a human disease mutation presented here have revealed a conserved structural motif that is critically important for kinesin-MT interactions. While prior studies have largely focused on the L11-α4-L12 region as the primary connection between kinesin and MTs, our results suggest the interaction is more extensive than previously appreciated, consistent with recent cryo-EM data (47), and that structural elements adjacent to this region, such as the 3_10_-helical element, may provide allosteric coupling critically important for the interaction. Our results further reveal that the presence of L12 is not sufficient for its function, lending support to the idea that a specific conformation of L12 couples to the kinesin enzymatic cycle.

## Materials and Methods

### C. elegans Methods

*C. elegans* were maintained on *E*.*coli* strain OP50 on Nematode growth medium (NGM) agar plates at 20 °C. *Punc-104*::*human Kif1a* vectors were prepared as described (14). 4ng of *Punc-104::human Kif1a* and 50 ng of *Podr-1::gfp* vectors were injected to *unc-104(e1265)* gonads by standard procedures (Mello et al., 1991). At the F1 generation, worms with visible GFP signal in head neurons were collected. Stable transformation was confirmed by transmission of extrachromosomal arrays to the F2 generation. Functional complementation of *unc-104* by human *KIF1A* was assessed by the motility of worms on NGM plates. Worms with visible markers were transferred to new plates and movement was recorded under an SteREO Discovery.V12 dissection microscope (Carl Zeiss, Jena, Germany) equipped with Orca-Flash 2.8 CMOS camera (Hamamatsu Photonics, Hamamatsu, Japan). Data were analyzed using Prism ver.7.

### Recombinant human KIF1A Assembly and Preparation

All KIF1A constructs were cloned into pAcebac1 vectors (Geneva Biotech, Genève, Switzerland) by Gibson assembly. cDNA encoding human *KIF1A* (Genbank: AB290172.1) was amplified by PCR. A DNA fragment encoding the mScarlet-2xStrepII tag was synthesized by gBlocks (Integrated DNA Technologies, Coralville, IA, USA). Full vector sequence was confirmed by Sanger sequencing. Truncated KIF1A constructs containing only the motor domain (M1-D393) were made from the amplified KIF1A gene. P305L mutations were introduced by PCR-based mutagenesis to both full-length and truncated constructs using Q5 plus DNA polymerase. Constructs containing the P305L mutation encoded a C-terminal sfGFP-2xStrepII tag. A bacmid was generated by transforming DH10EmBacY (Geneva Biotech) for all constructs. Insect Sf9 cells were maintained as a shaking culture in Sf-900II serum-free medium (SFM) (Thermo Fisher Scientific), or alternatively in ESF media (Expression Systems) at 27°C. To prepare baculovirus, 1 x 106 SF9 cells were transferred to each well of a 6-well plate. SF9 cells were transformed with bacmid DNA by Cellfectin (Thermo Fisher Scientific). The resulting baculovirus was amplified and the P2 cell culture medium was used for protein expression. To prepare recombinant proteins, 400 ml (2 x 10^6^ cells/ml) of cells were inoculated with P2 baculovirus stocks at a dilution of 1:100 and cultured for ∼65 hours at 27°C. Cells were harvested by centrifugation at 3000 × g for 5 min and frozen in liquid nitrogen. Frozen cells were stored at −80°C until the purification step.

### Recombinant human KIF5B Assembly and Preparation

All KIF5B constructs were cloned into pET28a vectors using Gibson assembly. A codon-optimized DNA sequence encoding the motor domain of human KIF5B (residues 1 – 420, Uniprot P33176) was used. A DNA fragment encoding the mScarlet-2xStrepII tag was synthesized by gBlocks (Integrated DNA Technologies, Coralville, IA, USA). Full vector sequence was confirmed by Sanger sequencing. The P276L mutation was introduced by PCR-based mutagenesis to both full-length and truncated constructs using Q5 plus DNA polymerase. KIF5B constructs were transformed into BL21-CodonPlus (DE3)-RIPLcells (Agilent) for protein expression. Cells were inoculated with 1:500 of overnight culture and were grown at 37°C in 1.5 L of LB media until optimal density at 600 nm (OD_600_) of 0.6. Cells were then induced with 0.1 mM isopropyl-β-D-thiogalactoside overnight (∼16 hours) at 18°C. Cells were harvested by centrifugation as 3000 × g for 15 min and frozen in liquid nitrogen. Frozen cells were stored at −80°C until the purification step.

### Purification of human KIF1A and KIF5B

For purification, 40 ml purification buffer (50 mM Tris, pH 8.0, 150 mM KCH3COO, 2 mM MgSO4, 1 mM EGTA, 10% glycerol) supplemented with 0.1% Triton-X100, 1mM ATP, 1 mM DTT, 1 mM PMSF and protease inhibitor mix (Promega) was used to resuspend frozen pellet that was thawed on ice. Cells were homogenized with several strokes in a dounce homogenizer and lysed by passing through an Emulsiflex C-3 (Avestin). Soluble lysate was obtained by centrifugation at 16000 × g for 20 min at 4°C. The lysate was mixed with 2 mL Strep-Tactin XT resin (IBA Lifesciences, Göttingen, Germany). The resin was washed extensively with wash buffer (50 mM Tris, pH 8.0, 450 mM KCH3COO, 2 mM MgSO4, 1 mM EGTA, 10% glycerol). Then, protein was eluted with elution buffer (50 mM Tris, pH 8.0, 150 mM KCH3COO, 2 mM MgSO4, 1 mM EGTA, 10% glycerol, 100 mM biotin). Eluted protein was concentrated to ∼ 500uL using Amicon Ultra 5 centrifugal filters (Merck, Darmstadt, Germany). The affinity purified protein was further separated by gel filtration using a Phenomenex Yarra 3µm SEC-4000 300 x 7.8mm column (Phenomenex, Torrance CA) in GF150 buffer (25 mM HEPES pH 7.4, 150 mM KCl, 1mM MgCl_2_) on an NGC Chromatography system (Bio-Rad Laboratories, Hercules, CA, USA). Peak fractions were pooled, concentrated in an Amicon filter again and flash frozen in liquid nitrogen. Protein concentrations were assessed using a Nanodrop One (ThermoFisher) and given as the total amount of measured fluorophore.

### TIRF assays

Glass chambers were prepared by acid washing as previously described (http://labs.bio.unc.edu/Salmon/protocolscoverslippreps.html) and double-sided sticky tape. Chambers were first incubated with 0.5 mg ml-1 PLL-PEG-biotin (Surface Solutions) for 10 min, followed by 0.5 mg ml-1 streptavidin for 5 min. Microtubules were diluted in BRB80-T (80 mM PIPES, 1mM MgCl2, 1mM EGTA, 5 µM taxol, pH 6.8) buffer. Taxol-stabilized MTs were flowed into streptavidin adsorbed flow chambers and allowed to adhere for 5–10 min. Unbound MTs were washed away using SRP90 assay buffer (90 mM Hepes pH 7.6, 50 mM KCH3COO, 2 mM Mg(CH3COO)2, 1 mM EGTA, 10 % glycerol, 0.1 mg/ml biotin–BSA, 0.2 mg/ml K-casein, 0.5 % Pluronic F127). Purified motor protein was diluted to indicated concentrations in the assay buffer with 2 mM ATP, and an oxygen scavenging system composed of PCA/PCD/Trolox. Then, the solution was flowed into the glass chamber. Images were acquired using a Micromanager software-controlled Nikon TE microscope (1.49 NA, 100× objective) equipped with a TIRF illuminator and Andor iXon CCD EM camera. Data were analyzed manually using ImageJ (FIJI) and statistical tests were performed in GraphPad PRISM 7.0c.

### ATPase Assays

MT-activated ATPase activity of KIF1A was analyzed in a NADH-coupled enzymatic assay (Huang et al., 1994). Initial rates of ATP hydrolysis by tail-truncated KIF1A or KIF1A^P305L^ at 37°C were measured as follows: KIF1A or KIF1A^P305L^ were diluted to 100nM in SRP90 supplemented with 2mM ATP, 0.01% Triton X-100, 1mM DTT, and 0.1mg/ml BSA. Coupled NADH oxidation system in the final reaction mixture was 0.1 mM NADH (Roche Diagnostics), 2 mM phosphoenolpyruvate (Sigma-Aldrich), 0.01 U pyruvate kinase (Sigma-Aldrich), and 0.03 U lactate dehydrogenase (Sigma-Aldrich). For each day of assays, taxol-stabilized MTs were assembled at 37°C for 30 min. and spun down by centrifugation at 150,000xg for 15 min. through 25% sucrose cushion. The MT pellet was resuspended in SRP90, and 0-10 µM MTs were added to the reaction. The reaction mixture was pre-incubated for 10 seconds at 37°C and the absorption at 340 nm was recorded in Eppendorf BioSpectrometer Kinetic. Data (n = 2-5) were fit to the Michaelis-Menten equation by nonlinear regression to calculate K_m_^MT^ and k_cat_ values using Prism.

### Optical tweezers assay

Slides, MTs, and polystyrene trapping beads were prepared as described previously (Rao et al., 2019). Briefly, carboxylate polystyrene beads (0.52 µm, Polyscience #09836-15) were coated with an Anti-Strep-tagII antibody (Abcam #ab76949) and α-casein. Glass coverslips (Zeiss #474030-9000-000) were cleaned with 25% HNO_3_ and 2 M NaOH, washed with ddH_2_O, air dried, and stored at 4°C. The flow cell was assembled with a glass slide, parafilm stripes, and a cleaned coverslip as described (Rao et al., 2019). Biotinylated MTs were attached to the coverslip surface via α-casein-biotin and streptavidin. The trapping assay was conducted as described (Rao et al., 2019)before except that HME60K50 buffer was used for all experiments (60 mM HEPES, 50 mM KAc, 2 mM MgCl2, 1 mM EGTA, 10% glycerol, 0.5% (w/v) Pluronic F-127). All force measurements were performed with a LUMICKS C-Trap®. The trap stiffness was set to 0.04-0.06 pN/nm, and the percentage of moving beads was between 10–45%, ensuring experiments at the single-molecule level (Brenner et al., 2020). A custom-made MATLAB program was used to analyze the data. Graphs were generated using Prism (GraphPad, version 8). This is an equation line. Type the equation in the equation editor field, then put the number of the equation in the brackets at right. The equation line is a one-row table, it allows you to both center the equation and have a right-justified reference, as found in most journals.

## End Matter

### Author Contributions and Notes

R.J.M and A.G. designed research and secured research funding, A.J.L., L.R., Y.A., K.C., K.O. performed research, D.W.N. conducted comparative analysis of published kinesin structures and created kinesin pseudo-models to aid in conceptualization. R.J.M, A.J.L., K.C., D.W.N, Y.A., L.R. analyzed data; and R.J.M, D.W.N and A.G wrote the paper. All authors edited the paper. Partial funding for this work was received from KIF1A.org. The authors declare no conflict of interest.

## Acknowledgments

The authors thank all the members of the MOM lab for their continual input and feedback on this project. The authors acknowledge the generosity and enthusiastic support of KIF1A.org and all the families suffering with KAND. This paper is dedicated to the Rosen family, and in particular to Susannah. We stand with you and all the KIF1A families! This work was generously supported by funds from KIF1A.org. The authors also thank Lia Boyle and Wendy Chung for helpful discussions on disease severity of the P305L mutation. The work was further supported by grants from NIGMS GM124889 (to R.J.M) and R01NS114636 (to L.R. and A.G.), The Japan Society for the Promotion of Science 20H03247, 19H04738, and 16H06536 (to S.N.). Hayaishi Memorial Scholarship for Study Abroad (to K.C.), and a JSPS Overseas Research Fellowship (to K.C.).

## References

1. N. Hirokawa, Y. Noda, Y. Tanaka, S. Niwa, Kinesin superfamily motor proteins and intracellular transport. Nat Rev Mol Cell Biol 10, 682–696 (2009).

2. Y. Okada, H. Yamazaki, Y. Sekine-Aizawa, N. Hirokawa, The neuron-specific kinesin superfamily protein KIF1A is a unique monomeric motor for anterograde axonal transport of synaptic vesicle precursors. Cell 81, 769–780 (1995).

3. V. Soppina, K. J. Verhey, The family-specific K-loop influences the microtubule on-rate but not the superprocessivity of kinesin-3 motors. Mol Biol Cell 25, 2161–2170 (2014).

4. V. Soppina et al., Dimerization of mammalian kinesin-3 motors results in superprocessive motion. Proc Natl Acad Sci U S A 111, 5562–5567 (2014).

5. J. W. Hammond et al., Mammalian Kinesin-3 motors are dimeric in vivo and move by processive motility upon release of autoinhibition. PLoS Biol 7, e72 (2009).

6. Y. Okada, N. Hirokawa, Mechanism of the single-headed processivity: diffusional anchoring between the K-loop of kinesin and the C terminus of tubulin. Proc Natl Acad Sci U S A 97, 640–645 (2000).

7. M. Tomishige, D. R. Klopfenstein, R. D. Vale, Conversion of Unc104/KIF1A kinesin into a processive motor after dimerization. Science 297, 2263–2267 (2002).

8. L. Boyle et al., Genotype and defects in microtubule-based motility correlate with clinical severity in KIF1A Associated Neurological Disorder. medRxiv 10.1101/2020.07.27.20162974, 2020.2007.2027.20162974 (2020).

9. M. Pennings et al., KIF1A variants are a frequent cause of autosomal dominant hereditary spastic paraplegia. Eur J Hum Genet 28, 40–49 (2020).

10. D. R. Gabrych, V. Z. Lau, S. Niwa, M. A. Silverman, Going Too Far Is the Same as Falling Short(dagger): Kinesin-3 Family Members in Hereditary Spastic Paraplegia. Front Cell Neurosci 13, 419 (2019).

11. E. Reid et al., A kinesin heavy chain (KIF5A) mutation in hereditary spastic paraplegia (SPG10). Am J Hum Genet 71, 1189–1194 (2002).

12. S. Esmaeeli Nieh et al., De novo mutations in KIF1A cause progressive encephalopathy and brain atrophy. Ann Clin Transl Neurol 2, 623–635 (2015).

13. P. Guedes-Dias et al., Kinesin-3 Responds to Local Microtubule Dynamics to Target Synaptic Cargo Delivery to the Presynapse. Curr Biol 29, 268–282 e268 (2019).

14. K. Chiba et al., Disease-associated mutations hyperactivate KIF1A motility and anterograde axonal transport of synaptic vesicle precursors. Proc Natl Acad Sci U S A 116, 18429–18434 (2019).

15. S. Kaur et al., Expansion of the phenotypic spectrum of de novo missense variants in kinesin family member 1A (KIF1A). Human mutation 10.1002/humu.24079 (2020).

16. E. Lopez et al., Identification of two novel KIF5A mutations in hereditary spastic paraplegia associated with mild peripheral neuropathy. J Neurol Sci 358, 422–427 (2015).

17. P. Enkhbayar, K. Hikichi, M. Osaki, R. H. Kretsinger, N. Matsushima, 3(10)-helices in proteins are parahelices. Proteins 64, 691–699 (2006).

18. L. Pauling, R. B. Corey, Configuration of polypeptide chains. Nature 168, 550–551 (1951).

19. R. S. Vieira-Pires, J. H. Morais-Cabral, 3(10) helices in channels and other membrane proteins. J Gen Physiol 136, 585–592 (2010).

20. G. L. Millhauser, Views of helical peptides: a proposal for the position of 3(10)-helix along the thermodynamic folding pathway. Biochemistry 34, 3873–3877 (1995).

21. U. Gether et al., Structural instability of a constitutively active G protein-coupled receptor. Agonist-independent activation due to conformational flexibility. J Biol Chem 272, 2587–2590 (1997).

22. Y. Ri et al., The role of a conserved proline residue in mediating conformational changes associated with voltage gating of Cx32 gap junctions. Biophys J 76, 2887–2898 (1999).

23. G. Woehlke et al., Microtubule interaction site of the kinesin motor. Cell 90, 207–216 (1997).

24. A. Caballero Oteyza et al., Motor protein mutations cause a new form of hereditary spastic paraplegia. Neurology 82, 2007–2016 (2014).

25. M. Fichera et al., Evidence of kinesin heavy chain (KIF5A) involvement in pure hereditary spastic paraplegia. Neurology 63, 1108–1110 (2004).

26. C. Goizet et al., Complicated forms of autosomal dominant hereditary spastic paraplegia are frequent in SPG10. Human mutation 30, E376–385 (2009).

27. M. C. Alonso, J. van Damme, J. Vandekerckhove, R. A. Cross, Proteolytic mapping of kinesin/ncd-microtubule interface: nucleotide-dependent conformational changes in the loops L8 and L12. EMBO J 17, 945–951 (1998).

28. R. Nitta, M. Kikkawa, Y. Okada, N. Hirokawa, KIF1A alternately uses two loops to bind microtubules. Science 305, 678–683 (2004).

29. J. Kumar et al., The Caenorhabditis elegans Kinesin-3 motor UNC-104/KIF1A is degraded upon loss of specific binding to cargo. PLoS Genet 6, e1001200 (2010).

30. B. G. Budaitis et al., Pathogenic Mutations in the Kinesin-3 Motor KIF1A Diminish Force Generation and Movement Through Allosteric Mechanisms. bioRxiv 10.1101/2020.09.03.281576, 2020.2009.2003.281576 (2020).

31. F. J. Kull, E. P. Sablin, R. Lau, R. J. Fletterick, R. D. Vale, Crystal structure of the kinesin motor domain reveals a structural similarity to myosin. Nature 380, 550–555 (1996).

32. J. R. Lee et al., An intramolecular interaction between the FHA domain and a coiled coil negatively regulates the kinesin motor KIF1A. EMBO J 23, 1506–1515 (2004).

33. S. Niwa et al., Autoinhibition of a Neuronal Kinesin UNC-104/KIF1A Regulates the Size and Density of Synapses. Cell reports 16, 2129–2141 (2016).

34. K. M. Ori-McKenney, J. Xu, S. P. Gross, R. B. Vallee, A cytoplasmic dynein tail mutation impairs motor processivity. Nat Cell Biol 12, 1228–1234 (2010).

35. H. T. Hoang, M. A. Schlager, A. P. Carter, S. L. Bullock, DYNC1H1 mutations associated with neurological diseases compromise processivity of dynein-dynactin-cargo adaptor complexes. Proc Natl Acad Sci U S A 114, E1597–E1606 (2017).

36. B. Y. Monroy et al., A Combinatorial MAP Code Dictates Polarized Microtubule Transport. Dev Cell 53, 60–72 e64 (2020).

37. C. Leduc et al., Molecular crowding creates traffic jams of kinesin motors on microtubules. Proc Natl Acad Sci U S A 109, 6100–6105 (2012).

38. R. Nitta, Y. Okada, N. Hirokawa, Structural model for strain-dependent microtubule activation of Mg-ADP release from kinesin. Nat Struct Mol Biol 15, 1067–1075 (2008).

39. M. P. Nicholas, L. Rao, A. Gennerich, An improved optical tweezers assay for measuring the force generation of single kinesin molecules. Methods Mol Biol 1136, 171–246 (2014).

40. A. Gennerich, Optical Tweezers : Methods and Protocols, Methods in Molecular Biology (Humana Press, New York, 2017).

41. K. Svoboda, C. F. Schmidt, B. J. Schnapp, S. M. Block, Direct observation of kinesin stepping by optical trapping interferometry. Nature 365, 721–727 (1993).

42. E. Meyhofer, J. Howard, The force generated by a single kinesin molecule against an elastic load. Proc Natl Acad Sci U S A 92, 574–578 (1995).

43. H. Higuchi, E. Muto, Y. Inoue, T. Yanagida, Kinetics of force generation by single kinesin molecules activated by laser photolysis of caged ATP. Proc Natl Acad Sci U S A 94, 4395–4400 (1997).

44. H. W. Schroeder, 3rd et al., Force-dependent detachment of kinesin-2 biases track switching at cytoskeletal filament intersections. Biophys J 103, 48–58 (2012).

45. S. Brenner, F. Berger, L. Rao, M. P. Nicholas, A. Gennerich, Force production of human cytoplasmic dynein is limited by its processivity. Sci Adv 6, eaaz4295 (2020).

46. K. J. Verhey et al., Light chain-dependent regulation of Kinesin’s interaction with microtubules. J Cell Biol 143, 1053–1066. (1998).

47. J. Atherton et al., Conserved mechanisms of microtubule-stimulated ADP release, ATP binding, and force generation in transport kinesins. Elife 3, e03680 (2014).

